# *Bacillus*: driver of functional states in synthetic plant root bacterial communities

**DOI:** 10.1101/2025.03.30.646149

**Authors:** Gijs Selten, Ronnie de Jonge

**Author notes:** Corresponding author: Ronnie de Jonge.

## Abstract

Plant roots release root exudates to attract microbes that form root communities, which in turn promote plant health and growth. Root community assembly arises from millions of interactions between microbes and the plant, leading to robust and stable microbial networks. To manage the complexity of natural root microbiomes for research purposes, scientists have developed reductionist approaches using synthetic microbial inocula, known as SynComs. In recent years, an increasing number of studies employed SynComs to investigate root microbiome assembly and dynamics under various conditions or with specific plant mutants. These studies have identified bacterial traits linked to root competence, but if and how these traits shape root microbiome dynamics across conditions is not well understood. To explore whether bacterial trait selection follows recurrent patterns, we conducted a meta-analysis of nine SynCom studies involving plant roots. Surprisingly, we observed that root communities frequently assemble into two distinct functional states. Further analysis revealed that these states are characterized by differences in the abundance of *Bacilli*. We propose that these *Bacilli*-associated functional states are driven by microbial interactions such as quorum sensing and biofilm formation. Additionally, we show that host activities, including root exudation and immune responses, influence the functional state of the root microbiome. Whether natural root communities also organize into distinct functional states remains unclear, but the implications could be significant. Functional diversification within root communities may influence the success and effectiveness of plant-beneficial bioinoculants, particularly *Bacilli*-based inoculants. To optimize microbiome-driven plant benefits, a deeper understanding of the mechanisms underlying functional state differentiation in root microbiomes is needed.

## Introduction

Assembly of the root microbial community, or microbiome, results from numerous interactions among soil micro-organisms and the plant root. Plants exude a complex array of organic compounds that attract soil microbes, which utilize these compounds for their growth. Consequently, these microbes travel to the rhizosphere and colonize the root surface (Dotaniya & Meena, 2015). The established root communities can, in turn, positively affect the plant by enhancing its health and growth (Berendsen *et al*., 2012; Pieterse *et al*., 2014; Stringlis *et al*., 2018). Inoculations of plant-beneficial bacterial strains on plant roots consistently demonstrate positive effects on plant health and growth when cultivated in laboratory settings (Asari *et al*., 2016; Blake *et al*., 2021; Carlsson & Huss-Danell, 2003; Fan *et al*., 2017; He *et al*., 2022; Pieterse *et al*., 2021). However, several studies reported that laboratory-based plant-beneficial effects disappear in agricultural field trials, where plants grow in complex natural microbiome environments (Bardin *et al*., 2015; Blake *et al*., 2021; Maplestone & Campbell, 1989; Moreira & May De Mio, 2015; Xu *et al*., 2011). This highlights a significant challenge in developing plant-beneficial bioinoculants or leveraging the root microbiome for plant-beneficial purposes: our limited understanding of root microbiome dynamics and functioning (Parnell *et al*., 2016). For example, a bioinoculant might perform variably in different complex community contexts (Eckshtain-Levi *et al*., 2020; Hu *et al*., 2021; Poppeliers *et al*., 2023; Wang *et al*., 2021). To overcome this problem, we need to better understand root microbiome assembly, its dynamics, and the root colonization strategies of microbes within. A comprehensive catalogue of the microbial functions, and their interactions, present in the root microbiome can potentially reveal what and how plant-microbe interactions and/or microbe-microbe interactions contribute to root community assembly.

Currently, little is known about the microbial functions that exist in complex soil microbial communities. Sequencing microbial marker genes provides insight in the microbiome profile but cannot reveal the functional information of microbial genes and functions. Metagenome prediction tools such as PICRUSt2 and Tax4Fun have been developed to address this issue, though they have been found inaccurate for soil and root microbiome samples due to database limitations (Aßhauer *et al*., 2015; Douglas *et al*., 2020; Sun *et al*., 2020). Sequencing the full metagenome of these samples can provide functional information, but even with high coverage, it is not possible to discover all microbial functions in the root microbiome (Di Bella *et al*., 2013). Additionally, metagenome sequencing is expensive and is affected by host-derived contamination of the metagenomic reads (Jovel *et al*., 2022), leading researchers to seek alternative methods. In the last 10 years, researchers have turned to synthetic community (SynCom) reconstitution experiments, which involve inoculating plants with small to large mixes of laboratory-cultured bacteria (Vorholt *et al*., 2017). This method requires isolating microbial strains from the roots, cultivating pure cultures from these strains, and sequencing the strains to gain insight in their microbial functions (Bai *et al*., 2015; Levy *et al*., 2018; Loper *et al*., 2012). Once these efforts have been completed, researchers have full control over the SynCom composition, allowing them to include or exclude bacterial strains based on taxonomy, functionality, or other criteria (Vorholt *et al*., 2017). Additionally, SynCom reconstitution experiments enable causal observations, as microbiome compositional changes or plant phenotypic changes can be explained by underlying microbial genetics using genome-wide association studies, and experimentally validated using mutant strains or drop-out experiments (Vorholt *et al*., 2017).

In contrast to natural microbiome studies, SynCom-based root communities are assembled from a predefined mix of cultured bacteria, each introduced in equal proportions at the start. This controlled setup enables researchers to study root community assembly in a stepwise manner, uncovering fundamental patterns, similarities, and differences in community formation. Given that microbiome assembly is governed by deterministic principles (Finkel *et al*., 2017; Müller *et al*., 2016), we hypothesize that root communities exhibit functional similarity. If so, this could suggest that specific bacterial functions associated with root competence are consistently selected across different communities.

To test this hypothesis, we conducted a meta-analysis of nine studies that inoculated SynComs of varying complexity on different host plants, or that cultured SynComs with diverse plant-derived compounds. For each study, we quantified the abundances of SynCom members and analyzed their genomic sequences to identify bacterial functions associated with root colonization. Additionally, these studies examined root microbiomes across multiple plant genotypes and environmental conditions, providing insights into how plant traits and environmental factors influence community composition and bacterial functional dynamics. We show that root communities exhibit functional overlap despite being composed of different, taxonomically diverse bacterial strains. Remarkably, our analysis reveals that in each study root microbiomes consistently segregate into two distinct functional states, defined by the presence or absence of specific bacterial functions. We identify *Bacillus* strains as key drivers of this functional state separation and show that the plant’s immune system plays an important role in shifting the root microbiome towards a *Bacillus*-associated state or a non-*Bacillus*-associated functional state.

## Methods

### Overview of studies

To conduct a meta-analysis of published SynCom studies, the amplicon reads of nine different studies were downloaded from NCBI on August 3^rd^, 2022 (BioProject numbers shown in Table 1, extensive description in Table S1). Additionally, the genomes and the genomic reads of the SynCom isolates that were used in all studies were downloaded from either *At*-SPHERE (www.at-sphere.com), NCBI (Bioproject: PRJNA297942 (used in Durán *et al*. (2021), Hou *et al*. (2021), Schandry *et al*. (2021), Wippel *et al*. (2021), and Wolinska *et al*. (2021)), *Lj*-SPHERE (www.at-sphere.com) (used in Wippel *et al*. (2021)), or from the DOE Joint Genome institute (GOLD study ID: Gs0019718 (used in Finkel *et al*. (2019), Finkel *et al*. (2020), Lebeis *et al*. (2015), and Qi *et al*. (2022)).

**Table 1.** Datasets included in this meta-analysis and corresponding BioProject IDs.

| Study | BioProject number |
| --- | --- |
| Duran_2021 | PRJEB43139 |
| Finkel_2019 | PRJNA531340 |
| Finkel_2020 | PRJNA543313 |
| Hou_2021 | PRJEB40980 |
| Lebeis_2015 | PRJEB9723 |
| Qi_2021 | PRJNA720397 |
| Schandry_2021 | PRJEB42439 |
| Wippel_2021 | PRJEB37695 |
| Wolinska_2021 | PRJEB45520 |

### Isolate quantification

We quantified SynCom isolate abundances using SyFi (SynCom fingerprinting), which leverages assembled genomes and raw genomic sequencing reads to generate strain-specific *16S rRNA* amplicon fingerprints (Selten *et al*., 2025). These fingerprints serve as references for pseudoalignment-based quantification of metagenomic amplicon reads. Amplicon reads were first error-corrected, chimera-filtered, and trimmed using DADA2 (version 1.26.0) with default settings, adjusting trimming lengths based on read quality (Callahan *et al*., 2016) (Table S2). SyFi then retrieved strain-specific *16S rRNA* V3-V4 or V5-V7 fingerprints for each study (Table S3) and performed pseudoalignment with Salmon (version 1.4.0), applying a minimum sequence identity score threshold of 0.95 to balance sensitivity and specificity (Patro *et al*., 2017; Selten *et al*., 2025). Absolute read counts were normalized by strain-specific *16S rRNA* copy numbers, generating an isolate count table.

Since SyFi requires genomic reads of SynCom strains, which were unavailable for some studies (Supplementary data 1), we conducted a parallel ASV-based analysis. Here, DADA2-filtered reads were processed in Qiime2 (version 2019.7) to infer ASVs (Bolyen *et al*., 2019), which were then matched against the public SynCom genome sequences at 99% sequence identity using VSEARCH (Rognes *et al*., 2016). The resulting isolate counts were normalized using SyFi’s *16S rRNA* copy number estimates.

We compared SyFi-pseudoaligned read counts with Qiime2-VSEARCH matched reads (Table S4). Unless Qiime2-VSEARCH captured substantially more reads, we prioritized SyFi’s isolate count dataset. In summary, the Qiime2-VSEARCH method was used only for Lebeis_2015 and Finkel_2020.

### Functional composition

To study bacterial functions in the root microbiome, we first annotated SynCom genomes using PROKKA (version 1.14.6) and eggNOG-mapper (version 2.1.4-2) (Cantalapiedra *et al*., 2021; Seemann, 2014). From these annotations, we extracted KEGG orthologs (KOs), quantified their occurrences per SynCom member, and compiled a genomic KO data table (Supplementary data 4) (Kanehisa *et al*., 2016).

To estimate KO abundances in the microbiome, we applied PICRUSt2, a metagenome prediction algorithm that matches ASVs to bacterial genomes and infers their functional composition table (Douglas *et al*., 2020). Instead of relying on database-matched genomes, we used our genomic KO table and microbiome isolate count table as input for PICRUSt2. This approach ensured that functional predictions were directly linked to SynCom members rather than inferred from unrelated reference genomes. The *16S rRNA* copy number normalization step in PICRUSt2 was omitted, as the microbiome isolate count table was already normalized for *16S rRNA* copy number.

### Diversity analyses

To investigate the similarity of the taxonomic and functional compositions of microbiome samples across and within studies, the Jaccard, Bray-Curtis, and Unweighted UniFrac distances between samples were calculated. The Jaccard distances indicate compositional similarity based on the presence/absence of features, while the Bray Curtis-distances indicate compositional similarity based on feature abundances. The Unweighted UniFrac distances, in turn, indicate compositional similarity based on presence/absence of taxa and their evolutionary relationships. For the Jaccard distances, the genus and KO count tables were converted to relative abundance tables and processed in Qiime2 (version 2019.7) (Bolyen *et al*., 2019), using the beta-diversity plugin with the Jaccard metric to calculate the between-sample Jaccard distances. To calculate Bray-Curtis distances (dissimilarities), we first added a pseudocount to the genus and KO abundance tables before normalizing it to relative abundances, as the Qiime2 version used (version 2019.7) did not support Bray-Curtis calculations with zeros. The normalized table was then imported into Qiime2 and processed with the beta-diversity plugin to compute Bray-Curtis dissimilarities (Bolyen *et al*., 2019). The Unweighted UniFrac distances were processed in Qiime2 as well, using the relative abundance isolate table without a pseudocount and the isolates’ *16S rRNA* sequences as input.

To assess the significance of compositional differences between sample groups, we conducted PERMANOVA analyses using adonis2 from the vegan R package (version 2.6-4) (Oksanen *et al*., 2013) on Jaccard, Bray-Curtis, and Unweighted UniFrac distance matrices. PERMANOVA quantifies the proportion of compositional variation explained by metadata variables such as study, SynCom, or root compartment.

### Sparse Partial Least Squares Discriminant Analysis

To identify bacterial KOs or isolates that drive sample group separation in the PCoA space of Jaccard distances, we performed sparse Partial Least Squares Discriminant Analysis (hereafter sPLS-DA) using mixOmics (version 6.24.0) in R (Rohart *et al*., 2017). sPLS-DA is a supervised classification method that prioritizes features distinguishing two classes. We first optimized hyperparameters through a grid search, selecting the optimal number of principal components, dataset subsets, iterations, and subset sizes. Principal components correspond to PCoA dimensions, while dataset subsets define sample grouping and partitioning strategy. The tuned hyperparameters were then used to run sPLS-DA, generating mean contribution scores for each KO or isolate, highlighting their role in group separation.

### Network analysis

Co-occurrence of bacterial genera across the nine studies was assessed using network analysis. Total abundance values were CLR-transformed and analyzed with FastSpar (SparCC) (version 1.0.0) (Watts *et al*., 2019). Median correlations were computed from 1,000 iterations, and significance was evaluated using 1,000 bootstraps.

## Results

### Functional overlap among rhizobacteria and root microbiomes

To explore similarities and differences in root community assembly based on microbial functions, we investigated the bacterial functional composition in nine publicly available SynCom datasets. These studies inoculated complex SynComs (> 20 strains) on plant roots or root-derived compounds. The SynComs originated from different soils, were investigated under varied environmental conditions or plant genotypes, and exhibited taxonomic diversity at multiple levels (Table S1). The most pronounced taxonomic differences were observed between US-derived (Levy *et al*., 2018) and European-derived SynComs (Bai *et al*., 2015) (Finkel_2019, Finkel_2020, Lebeis_2015, and Qi_2021 versus Duran_2021, Hou_2021, Schandry_2021, Wippel_2021, and Wolinska_2021) (Figure S1). Despite substantial taxonomic variation, functional overlap was evident—particularly at higher taxonomic levels such as family level—as revealed by KEGG Ortholog (KO) profiles. Across all datasets, 55.5% of detected KOs (4,267 out of 7,688 KOs) were shared (Figure S2), highlighting a core functional repertoire among root-associated bacteria. This suggests a degree of functional similarity in root microbiomes, regardless of geographic or taxonomic differences.

To assess whether functional overlap is also evident in SynCom-inoculated plant roots, we quantified SynCom members and extrapolated isolate abundances to KO abundances using the generated KO profiles. When excluding Lebeis_2015, we observed slightly higher functional similarity compared to genus similarity in 8 out of 9 studies (Figure 1), as indicated by a marginally lower PERMANOVA R^2^ value, suggesting reduced dissimilarity in compositions (R^2^: 0.24 for KO versus 0.26 for genus). Including Lebeis_2015 in the dataset, however, resulted in a higher similarity in genus composition than in functional composition (R^2^: 0.35 for KO versus 0.34 for genus). This discrepancy is driven by a relatively large fraction of unique KO functions (283) present in the SynCom of Lebeis_2015, distinguishing its functional composition from the other nine studies (Figure S2). In Lebeis *et al*. (2015), SynCom members were selected based on their root competence and varied responses to the plant’s immune system, particularly focusing on salicylic acid (SA) signaling pathways. This selection approach differs from other SynCom studies that primarily choose bacteria based on their prevalence in the root microbiome. Despite these differences, PERMANOVA results indicate that root microbiome assembly varies between SynComs composed of different genera or originating from different soils when assessing functional compositional similarity.

**Figure 1.**
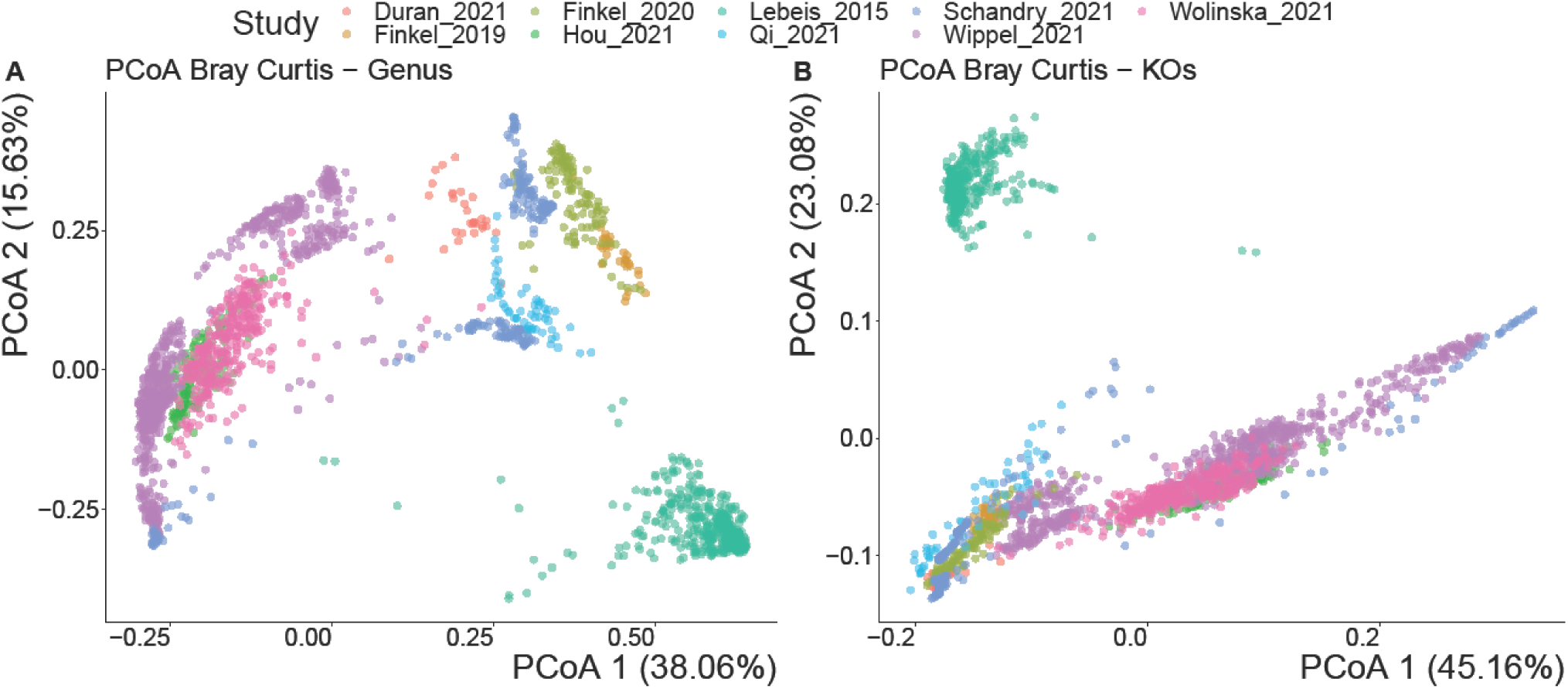
Principal coordinate analysis (PCoA) on Bray Curtis distances depicting the compositional differences among samples from the nine SynCom studies included in this meta-analysis. (A) PCoA based on Bray-Curtis distances of genus-level taxonomic profiles. (B) PCoA based on KO (KEGG Orthology) abundances. Each point represents an individual sample, with proximity indicating greater similarity in microbial composition.

### Root microbial communities assemble into two functional states

The limited overlap observed in the Bray Curtis analysis of the functional composition across studies (Figure 1) may stem from the metric’s sensitivity to conserved KOs present in multiple copies, such as ribosomal genes (Klappenbach, 2001; Louca *et al*., 2018). To address this, we employed Jaccard distances, which evaluated compositional similarity based on the presence or absence of genera or KOs, rather than their abundance (Figure 2). This approach revealed greater functional similarity compared to genus composition (R^2^: 0.37 for KO versus 0.43 for genus). The similarity became even more pronounced upon excluding the Lebeis_2015 dataset (R^2^: 0.24 for KO versus 0.35 for genus). These findings suggest that the bacterial genetic features associated with root competence are conserved across communities composed of different genera or derived from distinct soils.

**Figure 2.**
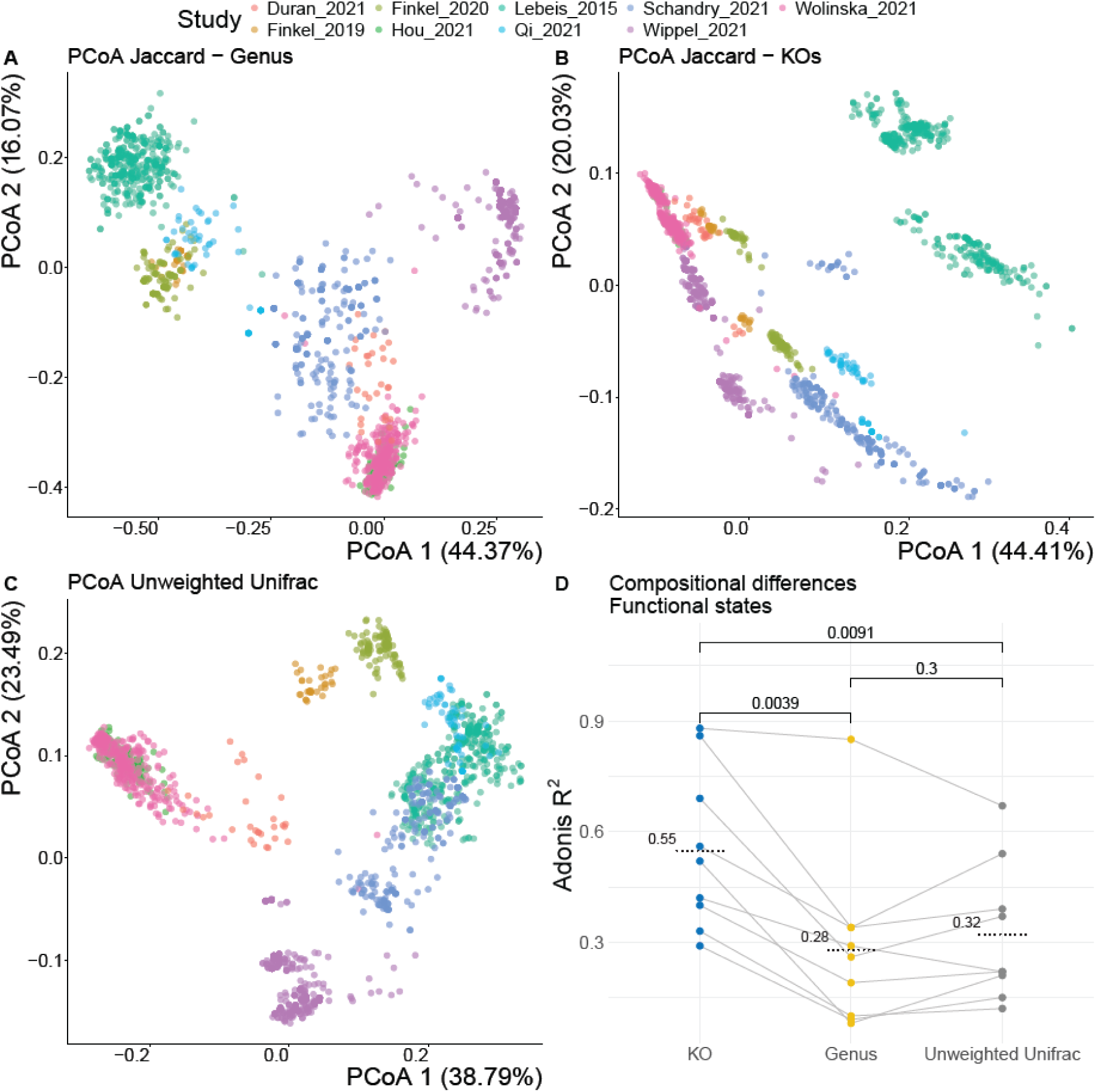
Principal Coordinate analysis (PCoA) plot based on Jaccard and Unweighted UniFrac distances. Jaccard distances between samples show compositional similarity between studies on the genus (A) and functional (KO) level (B). Unweighted UniFrac distances between samples indicate differential similarity between studies compared to the Jaccard distances (C). Compositional differences between the functional states that are evident in B are shown for the KO Jaccard distances, genus Jaccard distances, and Unweighted UniFrac distances (D). Significant differences between PERMANOVA R^2^ values in D are assessed by a paired Wilcoxon test and average values are indicated by dotted lines.

Remarkably, when examining Jaccard distances between functional root microbiomes within studies, we found that 7 out of 9 studies (all except Hou_2021 and Wolinska_2021) exhibit a division of samples into two distinct clusters or groups (Figures 2B, S3), a pattern not readily observed at the genus composition level (Figure 2A). To assess whether these two groups differ statistically, we applied K-means clustering (Hartigan & Wong, 1979) and analyzed their compositional differences. Both KO and genus compositions differed significantly between the two groups, though the differences were more pronounced at the functional level (mean R^2^: 0.28 for genus versus 0.55 for KO) (Figure 2D, Table S5). Given the clear divergence in functional composition with relatively minor taxonomic differences, we designate these groups as functional states—distinct microbiome configurations characterized by a specific a combination of functionalities. The fact that these functional states consistently emerge along the first two principal coordinate axes in nearly all studies (except for Schandry_2021; Figure S4), underscores their role as primary drivers of compositional differences. Although Hou_2021 and Wolinska_2021 do not exhibit a clear separation of groups (Figure S2), they still show substantial dissimilarity between the two K-means clusters (PERMANOVA R^2^ : 0.33 for Hou_2021 and 0.29 for Wolinska_2021). In conclusion, while it is widely recognized that deterministic principles guide root community assembly (Finkel *et al*., 2017; Lundberg *et al*., 2012; Müller *et al*., 2016), our findings reveal that these principles can lead to multiple functional outcomes, particularly evident in SynCom studies.

The observation that these two functional states are less pronounced yet still detectable at the genus level suggests a potential phylogenetic influence. To investigate this, we analyzed compositional differences between the functional states using Unweighted UniFrac distances, which consider both the presence or absence of taxa and their phylogenetic relationships (Figures 2C, S3). This analysis revealed clustering patterns like those observed at the genus level, with an unexpected overlap between Schandry_2021 and Lebeis_2015. Furthermore, the dissimilarity (R²) values between functional states were slightly higher when accounting for phylogeny (Figure 2D, Table S5), suggesting a modest phylogenetic signal associated with these distinct functional states.

### Bacillus is the main driver of distinct functional states

We have shown that the root microbiome organizes itself into two functional states, a pattern observed across nearly all studies, suggesting a common underlying mechanism. This may involve the presence or absence of specific bacterial functional groups, potentially originating from similar taxonomic lineages. Given that functional composition consistently accounts for the largest variation between these states, we first examined the contribution of individual KOs to the state separation across studies. To identify KOs associated with each state, we performed sparse Partial Least Squares Discriminant Analysis (sPLS-DA) on each study (Figures 3A, S5, Table S6). This supervised machine learning method extracts key features that distinguish different classes within the data.

**Figure 3.**
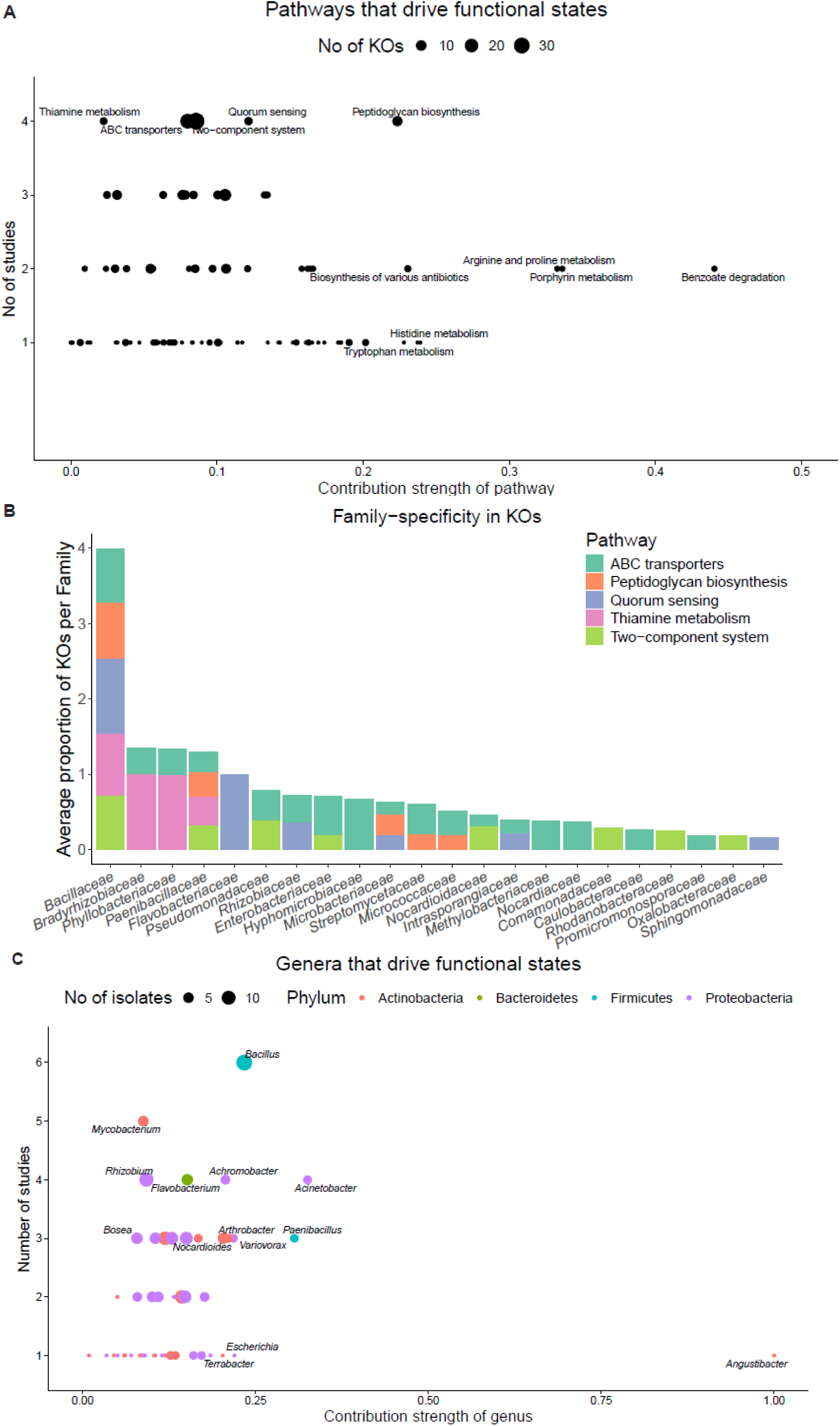
Contributions of pathways and genera to functional state separation. (A) The KOs that contribute to the functional state-separation were taken from the sPLS-DA analysis and grouped into the pathway they function in. The mean contribution strength of the pathway was calculated from these KOs (x-axis). The y-axis indicates the number of studies in which KOs of these pathways were found to have a contributing effect. The size of the coordinate indicates how many KOs have been found in that pathway. (B) For each of the top five pathways in A, the proportion of the contributing KOs belonging to different bacterial families was averaged. High proportions for a family-pathway combination indicate that contributing KOs can almost exclusively be found for that bacterial family. (C) The isolates that contribute to functional state-separation are grouped into genera displaying the mean contribution strength of the genus on the x-axis and the number of studies in which an isolate of the genus contributes to functional state-separation on the y-axis. The size of the coordinate indicates how many isolates of each genus contributes to functional state-separation, and the color indicates the phyla of the bacterial genera.

From the sPLS-DA, we identified 722 KOs including 669 unique ones, spanning 105 pathways that contribute to functional state separation across the nine studies. These KOs have a mean contribution of 0.082 (Table S6) and a summed contribution of 1-23.2 per study. Interestingly, in three studies (Finkel_2020, Qi_2021, and Schandry_2021), the functional states are distinguished by only one KO each (Figure S5), which are involved in peptidoglycan biosynthesis, biofilm formation, and intermembrane transport, respectively. This suggests that, in these cases, the presence or absence of a single KO is sufficient to predict functional states with high accuracy. In contrast, the number of KOs that contribute to functional state separation in three other studies is more than 200 (Figure S5), highlighting the significant variability in how functional states are separated across studies. The contributions of individual KOs to these functional states are often minimal, resulting in only slight differences in their relative abundances between functional states. The large number of contributing KOs across studies suggests that phylogenetically-related groups—whether comprising a few or many strains—tend to be present in one state and absent in the other. Since phylogenetically-related bacteria tend to be functionally similar, they often share near-identical KO profiles. Therefore, the absence of these groups in a given state results in the concurrent loss of multiple KOs, each contributing incrementally to functional state separation. To better interpret these patterns, we focus on the pathways most frequently associated with functional state separation: peptidoglycan biosynthesis, quorum sensing, two-component systems, ABC transporters, and thiamine metabolism (Figure 3A).

Across these pathways, 6.3 to 23.7% of KOs contribute to functional state separation (Table S7). Surprisingly, the most influential KOs originate from the conserved peptidoglycan biosynthesis pathway, which exhibits both the highest proportion of contributing KOs (9 out of 38; 23.7%) and the highest mean contribution (0.22) (Figure 3A). To further investigate their taxonomic origins, we filtered out widely distributed KOs, focusing instead on those that are more family-specific (Figure S6). Seven out of the nine peptidoglycan biosynthesis-related KOs meet this criterion, with six being restricted to *Bacillaceae* and *Paenibacillaceae* isolates (Figure S6). Many of the most influential KOs from the top five pathways are found in *Bacillaceae* isolates (Figure S6), with most being nearly exclusive to this family (Figure 3B). This pattern is especially pronounced in the peptidoglycan biosynthesis, quorum sensing, and thiamine metabolism pathways. These findings suggests that *Bacillaceae* strains may be the primary contributors to functional state separation across the nine SynCom studies.

To confirm whether *Bacillaceae* strains are the main drivers of the functional state separation, we performed sPLS-DA on the isolates for each study (Figures 3C, S7, Table S8). We identified 14 *Bacillus* strains in six out of nine studies, collectively contributing a mean of 0.31 to functional state separation, a substantial higher amount than any other bacterial genus (0.03-0.05).

Conclusively, we discovered that *Bacillaceae*-associated KOs and *Bacillus* strains drive the functional state separation (Figure 3). We observe that functional states do not align with genus-level compositional differences (Figures 2A, S3), despite being driven by *Bacillaceae* strains (Figures 3B, C). This suggests that the *Bacillaceae* clade may harbor many family-specific KOs, which collectively drive significant functional compositional differences. In turn, genus compositional differences are not greatly affected by *Bacillus* being a single entity. To confirm that *Bacillus* strains are indeed more present or abundant in one functional state compared to the other, we investigated the cumulative relative abundance of *Bacillus* strains in the functional states of each study (Figure S8). Apart from Qi_2021, which lacks *Bacillus* strains in the SynComs, all studies show a significantly higher proportion of *Bacillus* in one functional state compared to the other. Typically, one functional state is consistently depleted of *Bacilli*, while the other exhibits varying amounts of them. These findings suggest *Bacillus* establishment in the root microbiome follows an all-or-nothing strategy, though the ecological significance of *Bacillus* and this strategy remains unknown.

### Bacillus-associated states appear independent of the presence of other bacterial genera

The variable abundance of *Bacillus* in one functional state, paralleled to its absence or reduced presence in the other state, raises questions about the factors influencing the root microbiome’s functional state. One possible explanation is the effect of the other resident bacteria in the root microbiome, which may either positively or negatively affect *Bacillus* species or, on the other hand, thrive or decline because of the presence or absence of *Bacillus*. To test this hypothesis, we conducted a network analysis at the genus level across the nine studies to investigate which bacterial genera significantly correlate with *Bacillus*, positively or negatively. We focused on the top 10 most contributing bacterial genera to functional state separation (Table S8) and examined how these top contributors are related to each other. Interestingly, we found *Bacillus* to be isolated from all other top contributors, with only weak but significant negative correlations with *Acidovorax*, *Pseudomonas*, and *Flavobacterium*, which positively co-occur (Figure S9). This finding aligns with the understanding that *Pseudomonas* and *Bacillus* are often considered competent root microbiome inhabitants, primarily interacting competitively (Lyng & Kovács, 2023). The lack of positively correlating strains suggests that *Bacillus* is relatively independent of the presence and/or activities of other bacteria. On the other hand, the high abundance of bacterial genera such as *Pseudomonas*, might be more often associated with a non-*Bacillus* functional state. Further investigation is needed to better understand the specific interactions that lead to these distinct functional states in the root microbiome.

### Functional states correlate negatively with complexity

Since these findings derive from SynCom studies that inoculated a community of lower complexity compared to the natural root microbiome, we investigated whether the existence of two functional states is complexity-dependent. Interestingly, we found a significant, negative correlation between the SynCom complexity and the compositional differences between the functional states (Figure S10). Functional states are, therefore, more evident in smaller SynComs compared to larger SynComs. Whether this trend persists in natural communities, which are inherently more complex, remains uncertain and warrants further experimental investigation.

### Bacillus strains benefit from plant’s defense systems and loss of receptors

In addition to examining the influence of bacterial genera and SynCom complexity on functional states, we investigated how plant genotype and environmental factors might shape these states. Both plant genotype and environmental conditions have been shown to significantly impact root microbiome composition (Lundberg *et al*., 2012), potentially steering it toward one functional state or another. The studies included in this analysis encompassed a variety of treatments and conditions across multiple plant genotypes. This diversity enabled us to assess whether specific treatment or environmental factors are associated with functional states. To identify such associations, we conducted binomial statistical tests for overrepresentation, as detailed in Figures 4 and S11.

**Figure 4.**
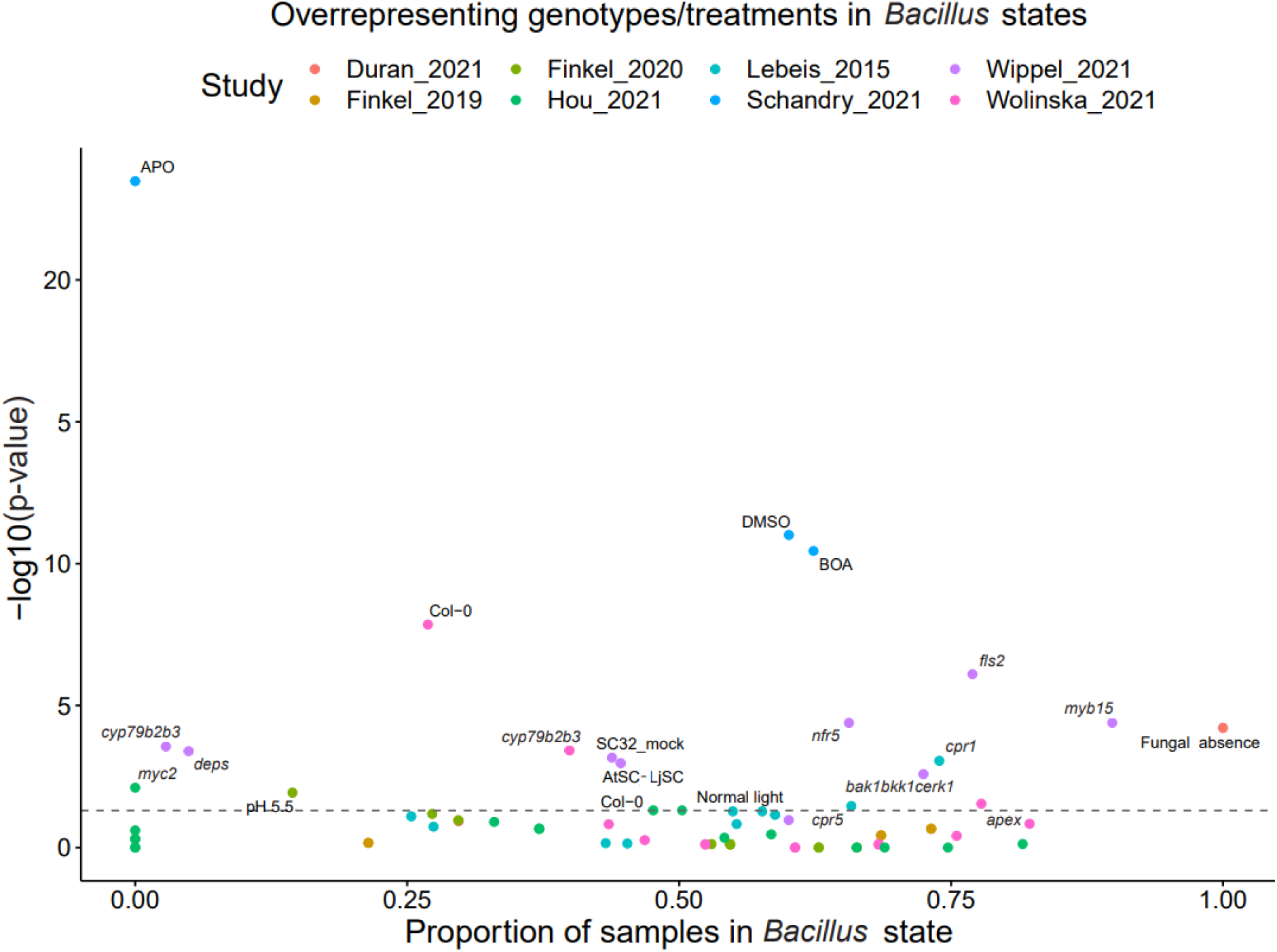
Overrepresentation analysis of genotypes and treatments in non-*Bacillus* or *Bacillus* functional states. Each point represents a unique genotype or treatment from one of the studies, color-coded by study for clarity. The x-axis denotes the proportion of samples within the *Bacillus* functional state, while the y-axis displays the negative logarithm (base 10) of the *p*-value from a binomial test, indicating the statistical significance of the association. A higher position on the y-axis signifies a stronger statistical association between the genotype or treatment and the functional state.

Our analysis revealed numerous significant associations between plant genotypes, treatments, and functional states within the root microbiome. For instance, bacterial communities were more likely to align with a *Bacillus*-associated functional state when the plant’s salicylic acid defense system was overexpressed (e.g., in *cpr1* and *cpr5* mutants) or when receptors for microbe-associated molecular patterns (MAMPs) were knocked out (e.g., *bak1bkk1cerk1*, *fls2*, *nfr5* and *apex* mutants) (Table S9). Conversely, impairing the plant’s defense system (e.g., in *deps*, *cyp79b2b3* and *myc2* mutants) was associated with a shift toward a non-*Bacillus*-associated functional state (Table S9). These findings suggest a complex interplay between plant immune responses and *Bacillus* strains, with different components of the immune system influencing these bacteria in varied ways. We hypothesize that locally-induced immunity via MAMP-triggered immunity (MTI) may suppress *Bacillus* strains, while systemic activation of plant immunity through mechanisms like induced systemic resistance (ISR) or systemic acquired resistance (SAR) may stimulate their presence. Additional findings indicate that factors such as benzoxazinoid APO, the presence of fungal pathogens, and lower pH levels negatively impact *Bacillus* strains (Figure 4). Based on these observations, it appears that environmental changes, aside from low pH, do not significantly affect functional states. However, comprehensive experimental validation is necessary to confirm these hypotheses.

## Discussion

SynCom reconstitution experiments have become increasingly popular for investigating the root microbiome (Marín *et al*., 2021; Martins *et al*., 2023; Shayanthan *et al*., 2022; Vorholt *et al*., 2017). These experimental set-ups provide insights into microbiome dynamics on a reduced scale while offering information on microbial functions within the root microbiome (Vorholt *et al*., 2017). However, the way root communities are established in SynCom reconstitution experiments differs from natural root communities. In natural soil, root communities develop from soil communities that vary spatiotemporally (Becker *et al*., 2022; Debray *et al*., 2022; Rüger *et al*., 2021). In contrast, SynCom experiments involve pooling bacterial species in proportionally similar abundances and inoculating them with plants to investigate root community establishment (Coker *et al*., 2022; Selten *et al*., 2024). The here-described meta-analysis demonstrates that inoculating taxonomically diverse bacterial strains results in functional overlap across studies, indicating a common functional similarity in bacterial root colonization strategies. This overlap, however, is not clearly visible on the abundance level of KOs, which may have multiple explanations. One possibility is that SynCom members exhibit a degree of soil-specificity in their functional profiles. Alternatively, the selection of rhizobacteria in some studies may not accurately represent the root microbiome, leading to differences in observed microbiomes. The way compositional differences are assessed—such as by using Bray-Curtis distances—may also influence results. KOs present in nearly all isolates, representing conserved bacterial functions, and those highly abundant within genomes tend to outweigh the effects of rare KOs in compositional similarity analyses. However, root-competence-associated functions can include both conserved and rare KOs (Levy *et al*., 2018; Selten *et al*., 2024).

Interestingly, we identified functional dissimilarity in the root microbiome that separates samples into two distinct states: a *Bacillus-*associated state and a non-*Bacillus*-associated state. The presence of multiple, dynamic, but stable root communities suggests the existence of plant enterotypes or rhizotypes—distinct compositional states in the root microbiome (Arumugam *et al*., 2011). In the human gut, enterotypes reach robust equilibria that can shift with prolonged changes in dietary patterns (Wu *et al*., 2011). Similarly, different enterotypes can be found in various human cultures due to dietary differences, analogous to how different rhizotypes may occur among different crop species (Berendsen *et al*., 2012). Since these distinct functional states are not found at the microbiome level where bacterial abundances are considered like they are in enterotypes, we believe the term “functional state” to be more appropriate than “rhizotype”.

The association between *Bacillus* species and the two functional states observed in root communities is complex and not yet fully understood. Both conserved and *Bacillaceae*-specific functions contribute to functional state separation, but their roles in determining the root microbiome’s overall composition remain speculative. *Bacillus* species employ quorum sensing and biofilm formation as key strategies for root colonization (Koua *et al*., 2020). Quorum sensing enables bacteria like *Bacillus* to coordinate collective behaviors, including biofilm formation, which enhance their ability to establish stable communities on plant roots. An in-depth analysis of biofilm formation within the *Bacillus* clade revealed considerable variation among biofilm formation strategies, underscoring its importance for *Bacillus* strain fitness (Abee *et al*., 2011; Cairns *et al*., 2014; Sun *et al*., 2022). Notably, *Bacillus* strains appear to cooperate in biofilm formation, which can be more effective than individual strategies (Kovács & Dragoš, 2019). This suggests that successful *Bacillus* biofilm establishment may promote a *Bacillus*-associated functional state. Conversely, failure to establish a biofilm could lead to exclusion from the root microbiome. This root colonization strategy, coupled with the fact that *Bacillus* is not necessarily the most abundant genus on the root, indicates that *Bacillus* abundance may be highly spatiotemporally organized, leveraging localized biofilm formation.

The *Bacillus* genus is widely utilized for plant-beneficial purposes, including pathogen biocontrol and plant growth promotion. Notable plant-beneficial *Bacillus* strains are *Bacillus amyloliquefaciens, Bacillus subtilis*, and *Bacillus velezensis* (Blake *et al*., 2021; Chowdhury *et al*., 2015; Gu *et al*., 2023; Kovács, 2019), although success rates in field trials was shown to be inconsistent (Blake *et al*., 2021). This variability may be attributed to the presence of both *Bacillus*-associated and non-*Bacillus*-associated functional states within natural root communities. These functional states could therefore significantly influence the efficacy of bioinoculants in agricultural applications, underscoring the necessity to investigate the factors that determine the establishment of *Bacillus*-associated functional states.

The plant’s immune system is a key factor in shaping its root communities (Teixeira *et al*., 2019). Activation of the immune system through mechanisms like MAMP-triggered immunity (MTI), systemic acquired resistance (SAR), or induced systemic resistance (ISR) can lead to various responses, such as reinforcement of cell walls, release of reactive oxygen species (ROS), and production of antimicrobial compounds (Belkhadir *et al*., 2014; Boller & Felix, 2009; Favaron *et al*., 2009; Ferrari *et al*., 2007; Galletti *et al*., 2008; Galletti *et al*., 2011; Jones & Dangl, 2006; Newman *et al*., 2013; Tsuji *et al*., 1992; Pieterse *et al*., 2014; Van Loon *et al*., 1998; Yu *et al*., 2022). These immune responses can affect root communities (Carvalhais *et al*., 2015; Doornbos *et al*., 2011; Lebeis *et al*., 2015; Pascale *et al*., 2020). Our findings show that the plant immune system can affect the functional state of the root community. The exact mechanisms driving this effect are, however, unclear, as both local and systemic components of the plant immune system appear to influence the functional state in different ways.

When the plant immune system is down-regulated, knocked down, or knocked out, we might expect a reduction in *Bacillus* populations due to the absence of defense responses that typically negatively affect competing gram-negative bacteria. These bacteria are reportedly more susceptible to diverse plant compounds, like salicylic acid, flavonoids, camalexins, and ROS (Adamczak *et al*., 2019; Rogers *et al*., 1996; Zhang *et al*., 2023), while gram-positive bacteria like *Bacillus* generally benefit from a thicker peptidoglycan cell wall, offering them better protection against such compounds (Zhang *et al*., 2023). *Bacillus* strains may exhibit greater tolerance to plant-derived antimicrobial metabolites compared to some other bacteria. However, they might still be adversely affected by root microbes that have also adapted to these metabolites. Conversely, systemic plant defenses appear to positively influence *Bacillus* strains. Given that the *Bacillus* genus harbors multiple plant-beneficial bacteria, they may positively respond to a plants’ cry for help following aboveground infection and subsequent activation of systemic immune responses (Blake *et al*., 2021; Chowdhury *et al*., 2015; Gu *et al*., 2023; Kovács, 2019). However, it remains unclear whether *Bacillus* species are predominantly plant-beneficial, commensal or pathogenic, as research on this genus may be subject to bias. Additionally, our findings suggest that lignins, which have been found to negatively affect *Bacillus* and other gram-positive species (Alzagameem *et al*., 2019; Guo *et al*., 2018), may explain the presence of a *Bacillus*-associated functional state in the lignin-deficient *myb15* mutant.

In addition to examining microbe-microbe and plant-microbe interactions, we evaluated the impact of various abiotic and biotic factors on functional states. Schandry *et al*. (2021) demonstrated that gram-negative bacteria exhibit higher tolerance to the benzoxazinoid APO compared to gram-positive bacteria, a result we corroborate (Figure 4). Notably, all APO samples were associated with the non-*Bacillus*-associated state. Durán *et al*. (2018) reported no significant difference in *Bacillus* abundances in the root microbiome with or without a fungal pathogen, although *Bacillus* levels were slightly higher in the absence of the pathogen, a trend supported by our analysis. Conversely, *Bacillus* bacteria are known to associate with beneficial fungi (Kjeldgaard *et al*., 2019; Long *et al*., 2017; Steffan *et al*., 2020; Toljander *et al*., 2006). However, experimental validation is required to confirm positive interactions with beneficial fungi and negative interactions with pathogenic fungi. Additionally, inorganic compounds also influence the functional state. We observed *Bacillus* depletion when plants were grown at pH 5.5, consistent with studies showing that *Bacillus* growth is severely inhibited at low pH (Valero *et al*., 2003; West *et al*., 1985). Overall, our findings are in line with previous research on environmental perturbations that significantly affect the functional state of root communities.

Finally, the presence of *Bacillus*-associated functional states in natural soils remains uncertain. Our analysis indicates that the presence of functional states correlates negatively with SynCom complexity. In SynComs with moderate complexity, functional states are less distinct compared to those in medium or low complexity settings, although differential *Bacillus* abundances are still observable. This negative correlation may be influenced by technical biases, as more complex communities tend to exhibit higher functional diversity, potentially diminishing the impact of *Bacillus*-derived functions on the overall community. Exploring *Bacillus* populations in natural rhizospheres through publicly available datasets could shed light on whether *Bacillus* strains drive functional state separation in natural root communities. Understanding the existence of multiple *Bacillus*-driven functional states in the root microbiome would significantly enhance our fundamental understanding of root microbiome dynamics and establishment. Such knowledge could improve bioinoculant development, as the functional state might affect the bioinoculant’s success in integrating into the root microbiome. Moreover, root microbiomes could potentially be engineered to favor specific functional states that support plant-beneficial microbes. Although many findings from this meta-analysis require experimental validation, the observed effects of these functional states on microbial functions at the presence/absence level highlight the potential of utilizing SynComs to study the root microbiome.

## Acknowledgments and funding

The authors want to thank Corné M.J. Pieterse for critical feedback on the manuscript. This study was supported by the Novo Nordisk Foundation Grant no. NNF19SA0059362.

## Competing interests

The authors declare no competing interests.

## Author contributions

G.S. and R.d.J. conceptualized the conducted meta-analysis. G.S. gathered and investigated the data, conducted the bioinformatic analyses, visualised the results, and wrote the manuscript. R.d.J. oversaw project administration, supervision, and revised the manuscript.

## Data availability

The supplementary tables and datasets are available at the following Zenodo link: https://doi.org/10.5281/zenodo.14982431. This repository includes an R script that generates all figures and supplemental figures in the manuscript and performs the sPLS-DA analysis. Additionally, a zip file containing all necessary files to run the R script is provided.

## Supplemental figures

**Figure S1.**
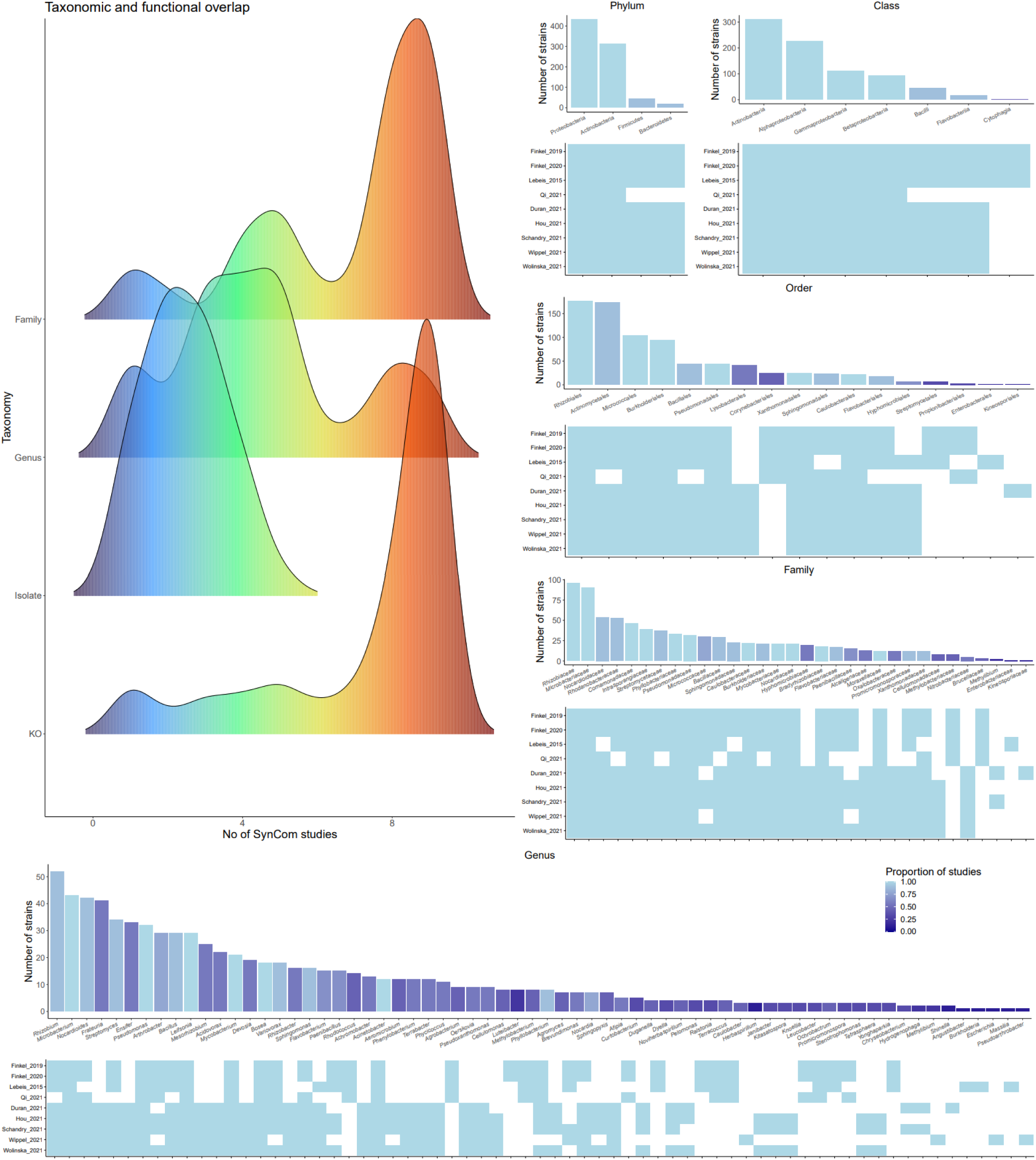
Taxonomy of SynCom members used in the nine SynCom studies included in this meta-analysis. Distribution plot of the number of unique taxonomic groups and KOs present per number of SynCom studies (x-axis) (top left). For each taxonomic level - phylum, class, order, family, and genus - the number of strains is displayed across all SynCom studies (y-axis), with color indicating the number of studies in which each taxonomic group is included (top-right and bottom). The heatmap below the bar charts indicates which SynCom study contains the specific taxonomic group.

**Figure S2.**
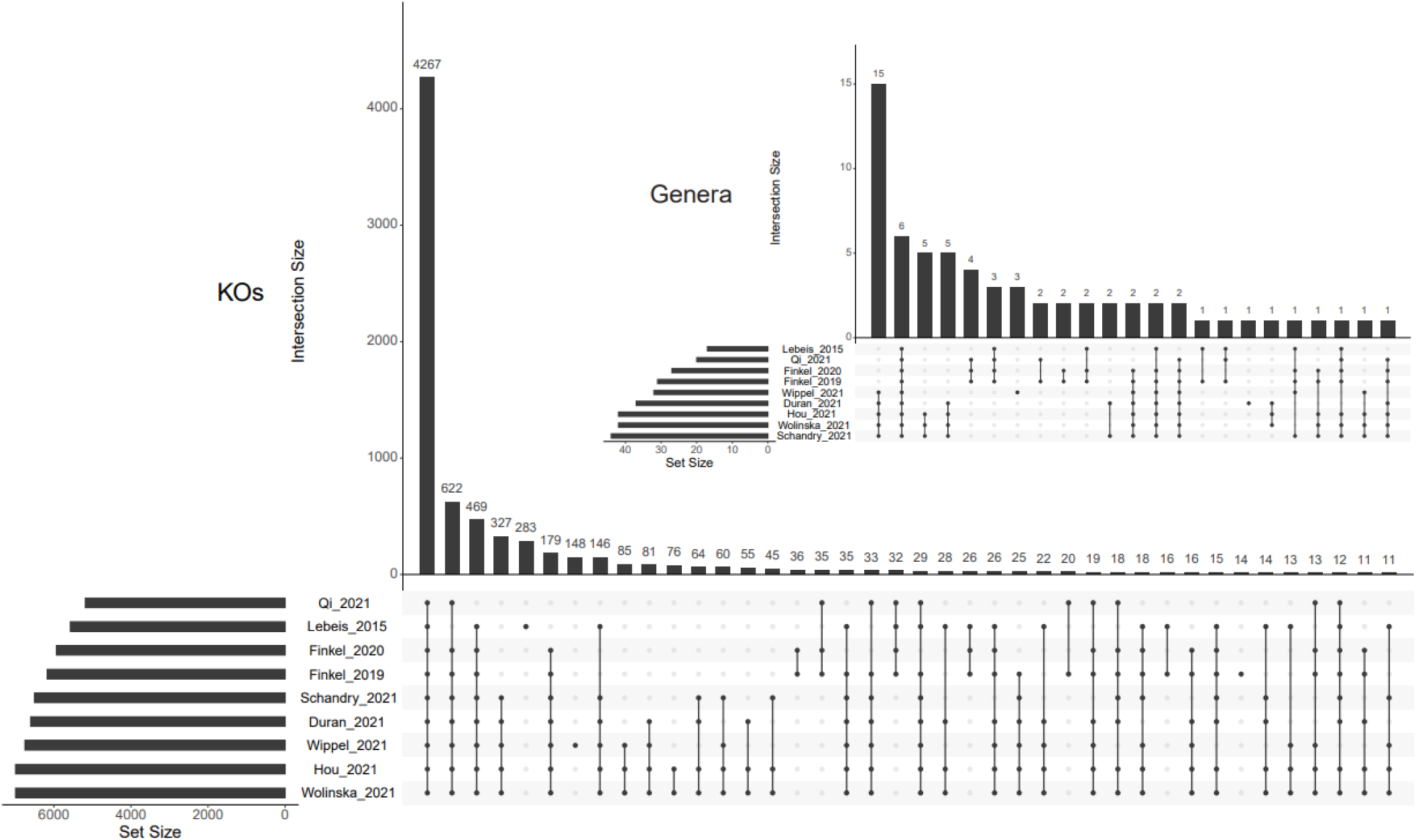
Overlap of KOs (bottom) and genera (top) that are present in the SynComs used in the nine studies. UpSet plot depicting the overlap in KOs and genera between the nine SynCom studies included in this meta-analysis.

**Figure S3.**
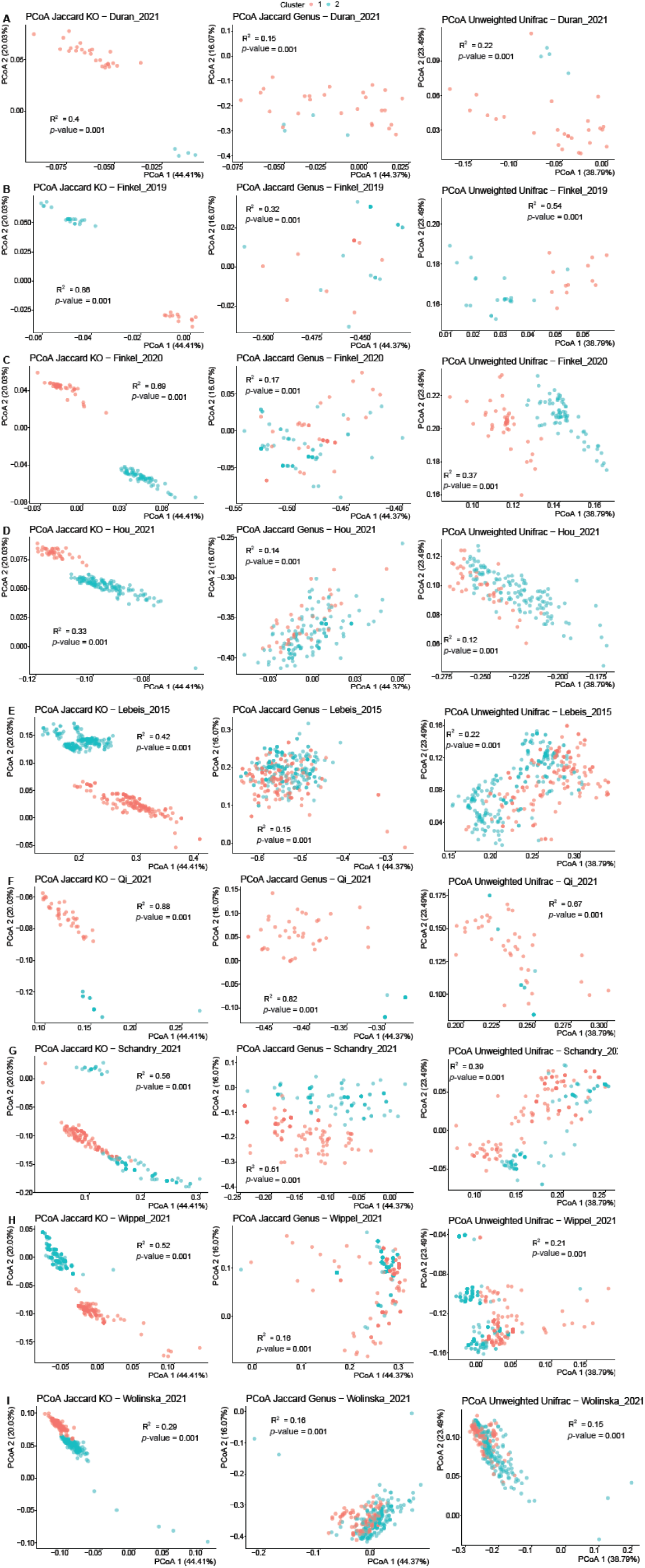
Principal Coordinate Analysis (PCoA) plots of KOs, genera and taxonomy of the nine SynCom studies included in this meta-analysis. KO composition based on Jaccard distances (left), genus composition based on Jaccard distances (middle), and taxonomic composition based on Unweighted UniFrac distances (right). Pe study, K-means clustering was used to group the samples into two main groups based on the KO composition which are indicated by the red and turquoise color. For the genus composition based on Jaccard distances PCoAs and the Unweighted UniFrac PCoAs, the cluster colors are based on the groups made according to the KO composition. For most studies there is a clear distinction in functional composition between the defined functional states, statistically assessed by an Adonis PERMANOVA.

**Figure S4.**
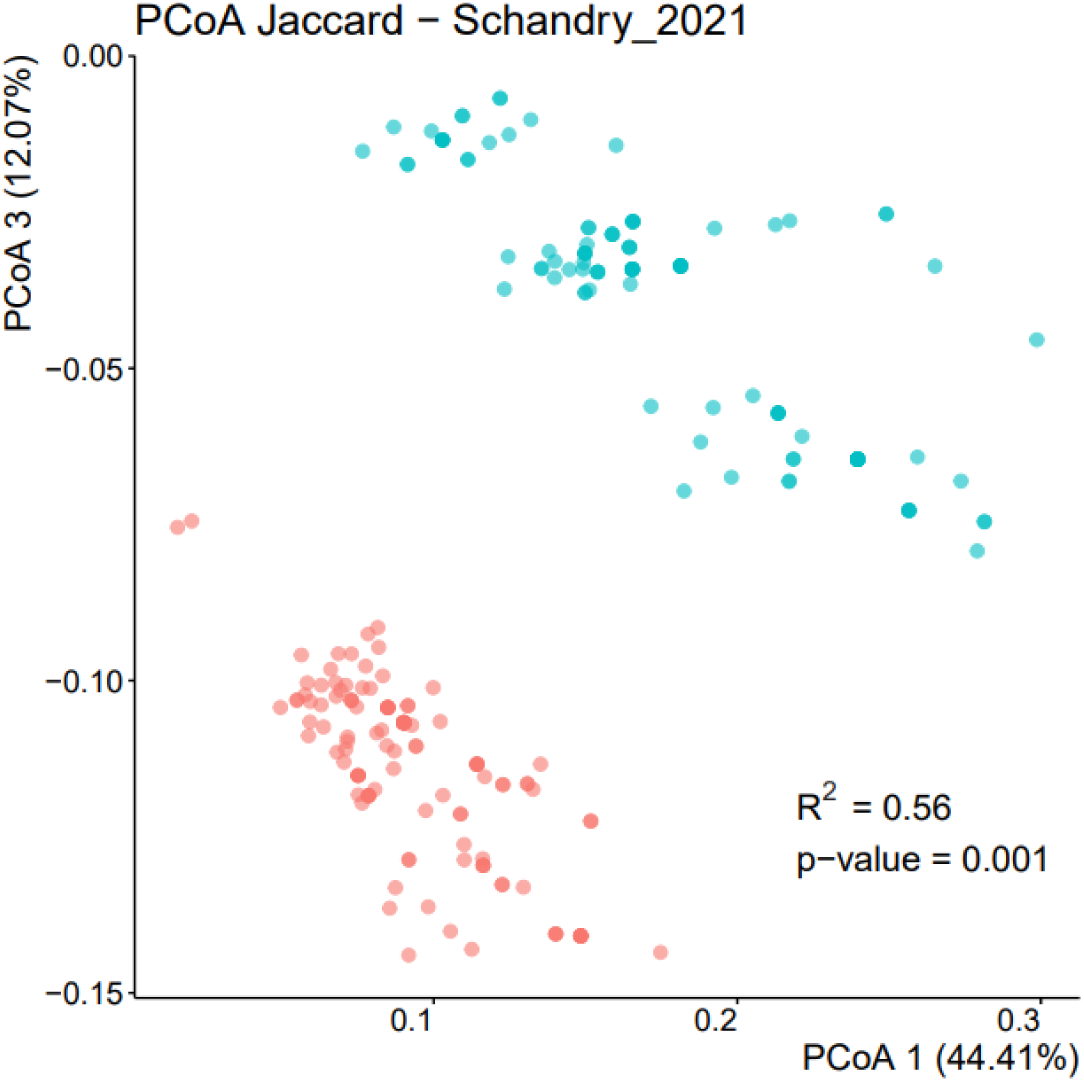
Principal Coordinate Analysis (PCoA) plot illustrating the functional states in Schandry_2021. PCoA on the KO composition based on Jaccard distances in Schandry_2021 on the first and third coordinate axes.

**Figure S5.**
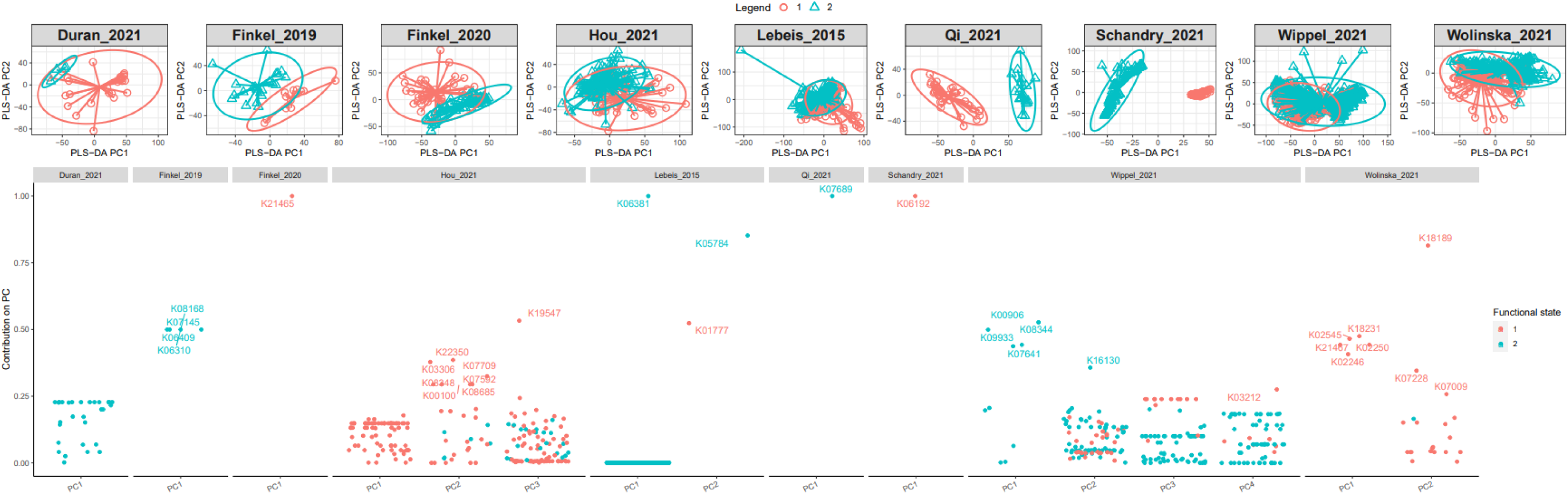
Contributions of KOs to functional state separation indicated by sPLS-DA analysis. sPLS-DA analysis was conducted on the two principal component axes that define distinct functional states (top). The analysis identifies the KOs that contribute most to the separation of functional states on each principal component (bottom; y-axis). The x-axis in the bottom panel indicates the principal component to which the KO contributes. Colors represent the functional states associated with each KO.

**Figure S6.**
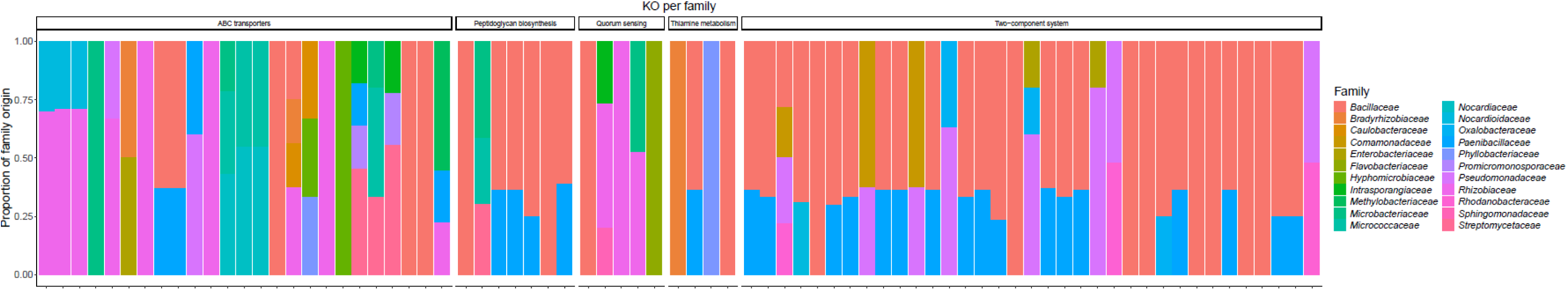
Familial origins of key KOs contributing to functional state separation. Stacked bar graph depicting the familial origin of the contributing KOs for the top five pathways (grouped and labeled in the top horizontal bar) that contribute most frequently to functional state separation across studies (Figure 3A). Included KOs are those with minimal proportion > 0.15 (full results in Table S7).

**Figure S7.**
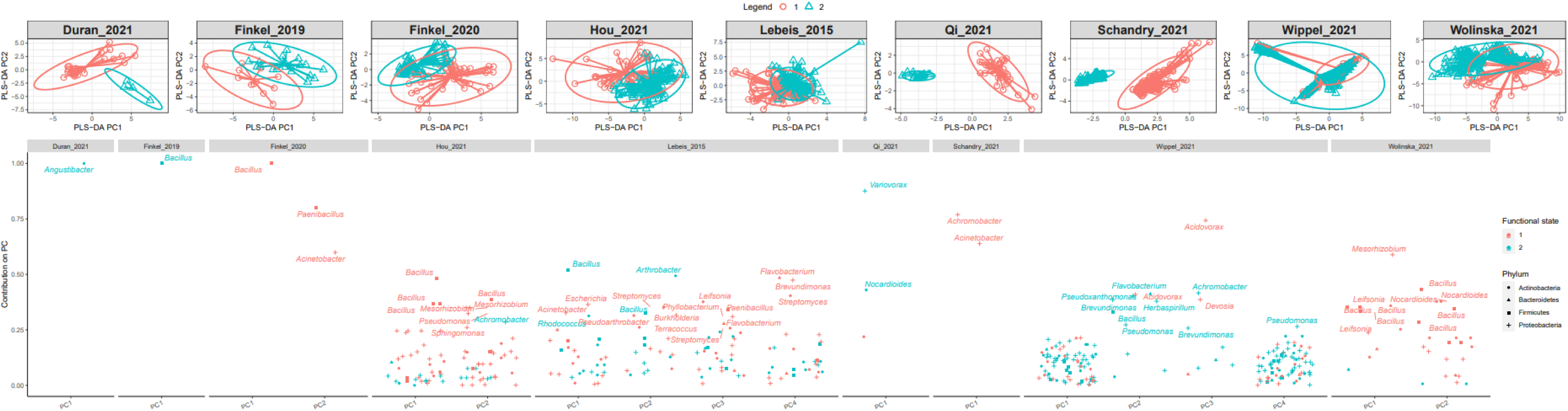
Contributions of genera to functional state separation indicated by sPLS-DA analysis. sPLS-DA analysis was conducted on the two principal component axes that define distinct functional states (top). The analysis identifies the isolates that contribute most to the functional state separation on each principal component (bottom; y-axis). The x-axis in the bottom panel indicates the principal component to which the isolate contributes. Colors represent the functional states associated with each isolate.

**Figure S8.**
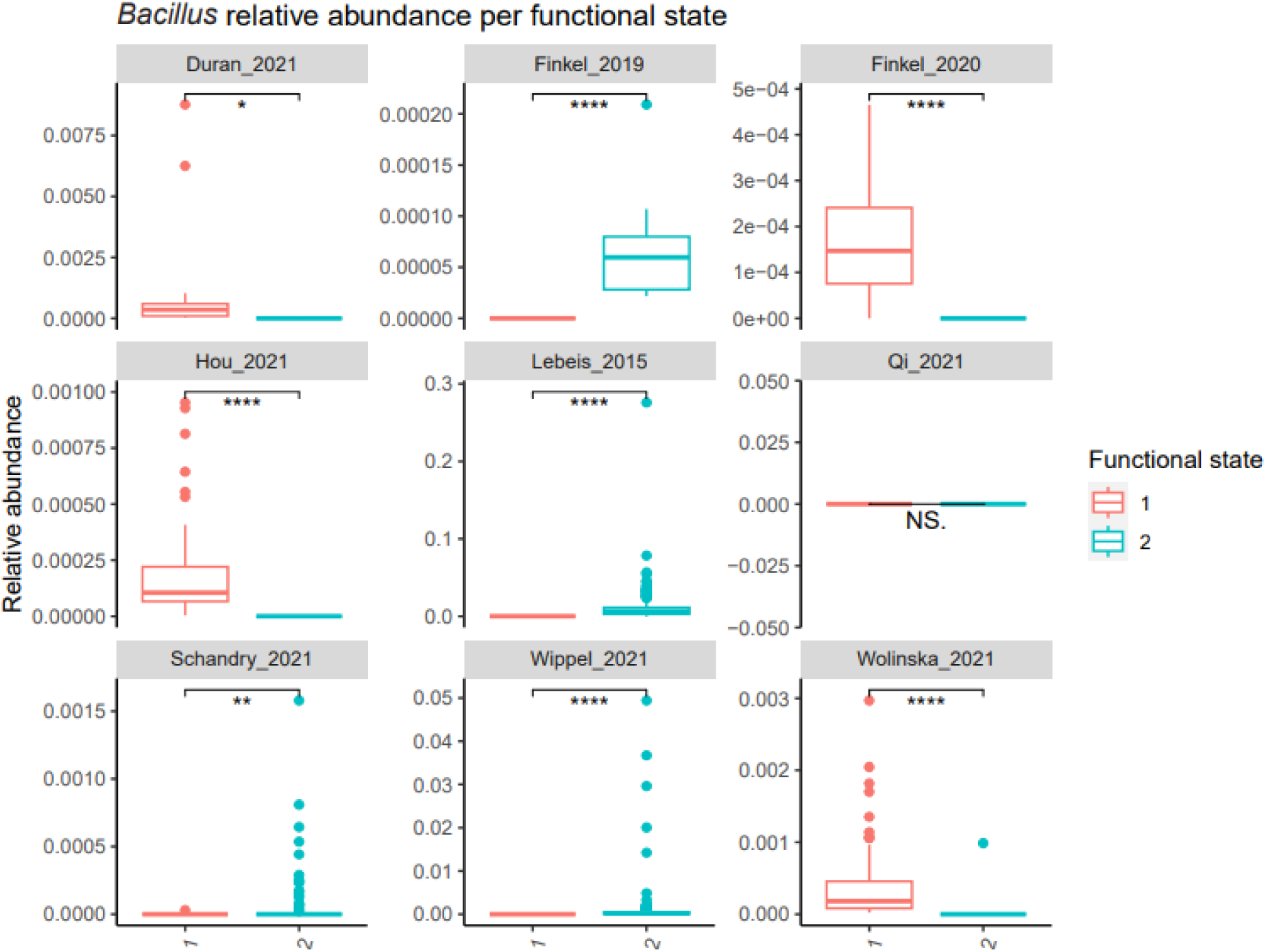
Relative abundance of *Bacillus* in the two functional states. Box plots describing the relative abundance of *Bacillus* in the two functional states across the nine studies included in this meta-analysis. Significant differences between functional states that derive from pairwise t-tests are indicated by * (*p* > 0.05 (ns), *p* < 0.05 (*), *p* < 0.01 (**), *p* < 0.001 (***), *p* < 0.00001 (****)).

**Figure S9.**
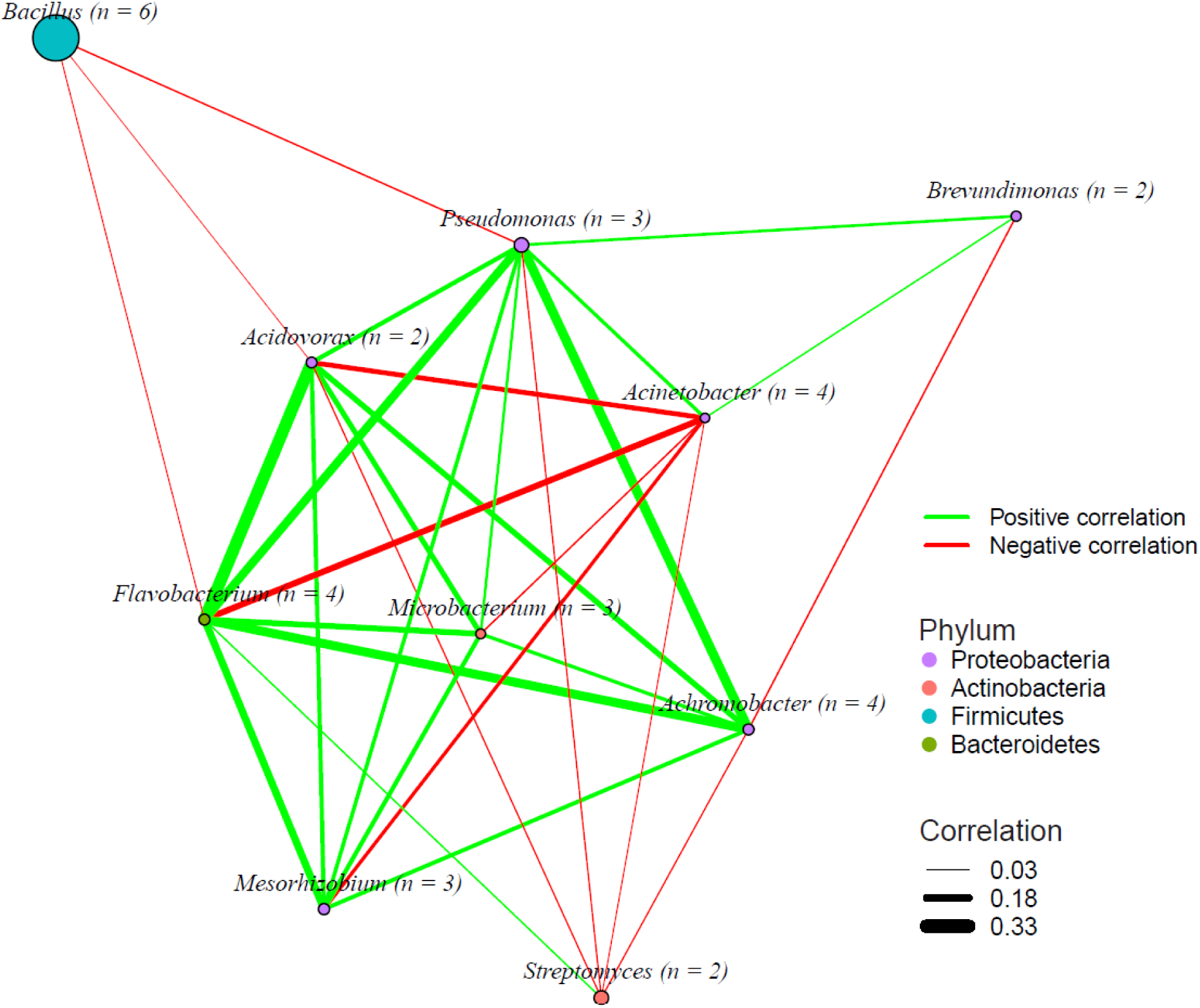
Network analysis at the genus level across the nine studies included in this meta-analysis. This network analysis illustrates microbial interactions at the genus level across nine studies. The size of each node represents its contribution to functional state separation, with *n* indicating the number of studies in which the bacterial genus contributed. The width of the edges indicates the correlation strength, with positive correlations shown in green and negative correlations in red. All displayed edges are statistically significant (*p* < 0.05) assessed by bootstrap resampling (*n* = 1000).

**Figure S10.**
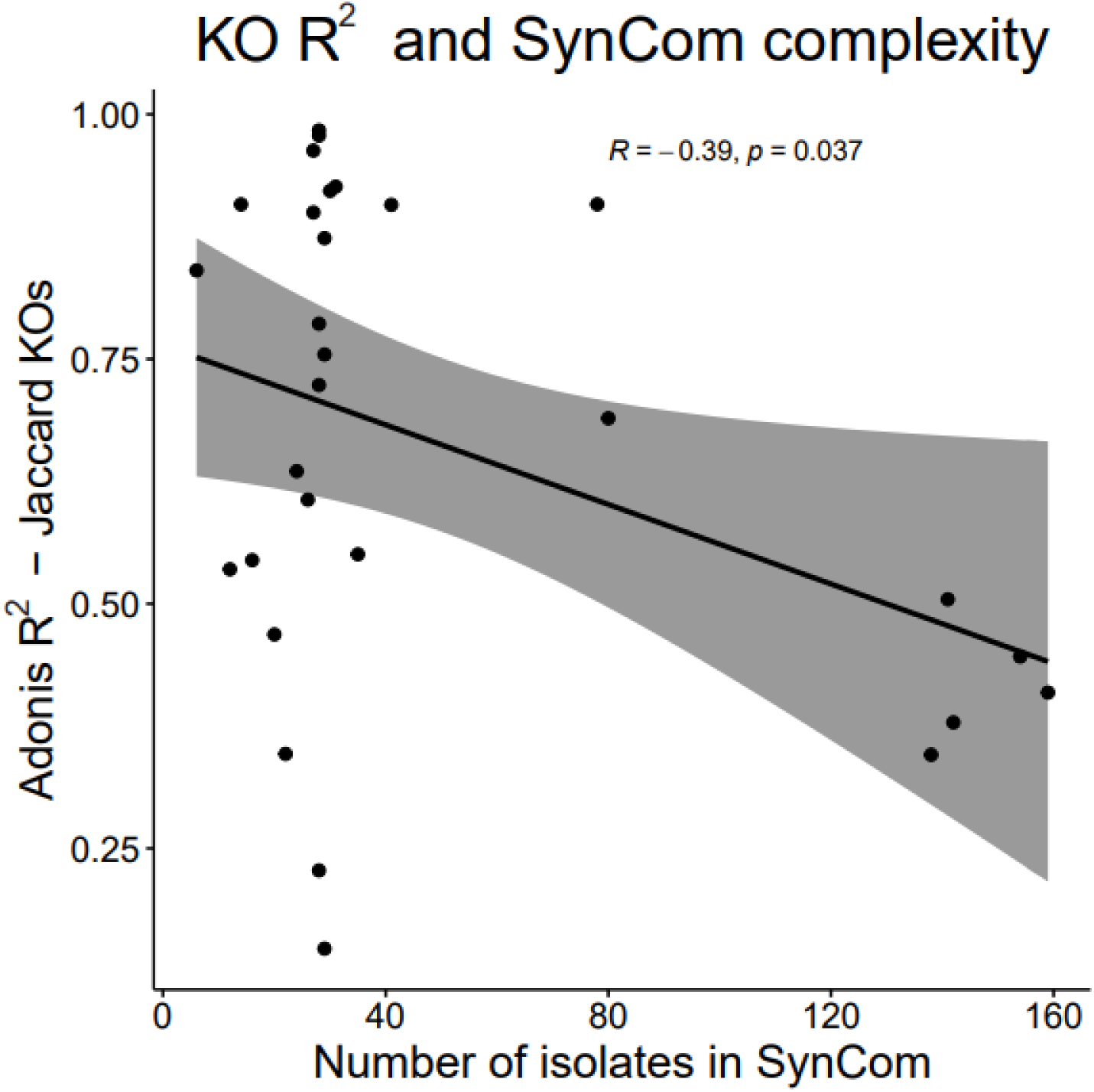
Correlation between compositional dissimilarity between functional states and SynCom complexity. Scatter plot depicting the inverse relationship between the number of isolates in the SynCom (x-axis) and the degree of separation between the functional states, as measured by the PERMANOVA R^2^ values of compositional differences (y-axis). Notably, the Qi_2021, Schandry_2021, and Wippel_2021 datasets included multiple SynComs (three, four, and ten, respectively), resulting in additional datapoints.

**Figure S11.**
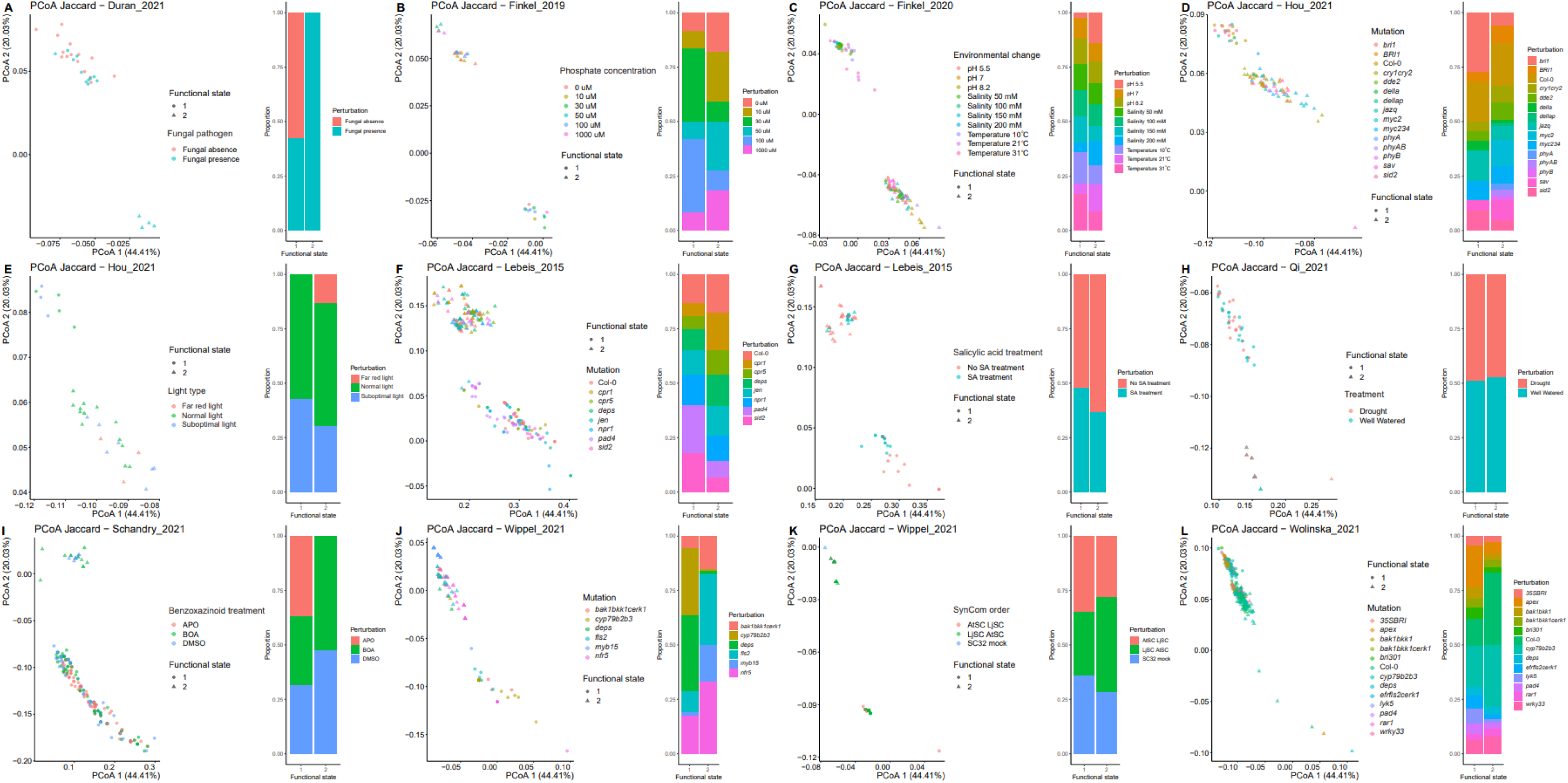
Principal Coordinate Analysis (PCoA) plots illustrating the distribution of sample treatments and host genotypes across two functional states. In each panel, samples are plotted based on their KO composition, with functional states differentiated by shape: circles represent functional state 1, and triangles denote functional state 2. Colors correspond to various treatments or host genotypes applied in each study, including Arabidopsis and Lotus (for Wippel *et al*., 2021). Bar plots adjacent to each PCoA plot display the distribution of treatments and genotypes within each functional state. For studies implementing combined treatments and genotypes (Hou_2021, Lebeis_2015, and Wippel_2021), separate plots are provided for treatments and genotypes. Detailed descriptions of treatments and host genotypes are available in Tables S1 and S9, respectively. In panel K, the ‘SynCom order’ indicates the sequence in which SynComs were inoculated onto the host, simulating the establishment of a root microbiome followed by invasion with another inoculum.

## Notes

https://doi.org/10.5281/zenodo.14982431

### Competing Interest Statement

The authors have declared no competing interest.

